# *Listeria monocytogenes* MenI encodes a DHNA-CoA thioesterase necessary for menaquinone biosynthesis, cytosolic survival, and virulence

**DOI:** 10.1101/2020.12.21.423900

**Authors:** Hans B. Smith, Tin Lok Li, Man Kit Liao, Grischa Y. Chen, Zhihong Guo, John-Demian Sauer

## Abstract

*Listeria monocytogenes* is a Gram-positive intracellular pathogen that is highly adapted to invade and replicate in the cytosol of eukaryotic cells. Intermediate metabolites in the menaquinone biosynthesis pathway are essential for the cytosolic survival and virulence of *L. monocytogenes*, independent of the production of MK and aerobic respiration. Determining which specific intermediate metabolite(s) are essential for cytosolic survival and virulence has been hindered by the lack of an identified DHNA-CoA thioesterase essential for converting DHNA-CoA to DHNA in the MK synthesis pathway. Using the recently identified *Escherichia coli* DHNA-CoA thioesterase as a query, homology sequence analysis revealed a single homolog in *L. monocytogenes*, LMRG_02730. Genetic deletion of LMRG_02730 resulted in an ablated membrane potential, indicative of a non-functional electron transport chain (ETC) and an inability to aerobically respire. Biochemical kinetic analysis of LMRG_02730 revealed strong activity towards DHNA-CoA, similar to its *E. coli* homolog, further demonstrating that LMRG_02730 is a DHNA-CoA thioesterase. Functional analyses *in vitro*, *ex vivo*, and *in vivo* using mutants directly downstream and upstream of LMRG_02730 revealed that DHNA-CoA is sufficient to facilitate *in vitro* growth in minimal media, intracellular replication, and plaque formation in fibroblasts. In contrast, protection against bacteriolysis in the cytosol of macrophages and tissue specific virulence *in vivo* requires the production of DHNA. Taken together, these data implicate LMRG_02730 (renamed MenI) as a DHNA-CoA thioesterase and suggest that while DHNA protects the bacteria from killing in the macrophage cytosol, DHNA-CoA is necessary for intracellular bacterial replication.

## INTRODUCTION

*Listeria monocytogenes* is a Gram-positive, intracellular pathogen capable of causing the severe disease listeriosis, resulting in an approximately 30% mortality rate^1^. To cause disease, *L. monocytogenes* must invade host cells and access their cytosol to establish its replicative niche^2^. *L. monocytogenes*, like other cytosolic pathogens, is remarkably well adapted to life within the eukaryotic cytosol^3^. In contrast, bacteria which are not specifically adapted to the restrictive cytosolic environment are unable to survive and replicate^4–8^. *L. monocytogenes* utilizes a myriad of virulence factors to facilitate entry into cells where it is initially encapsulated in a phagosome^2,9^. The pore-forming toxin listeriolysin O (LLO), encoded by the gene *hly,* facilitates escape of *L. monocytogenes* into the cytosol^10^. Infection is then disseminated to neighboring cells via the protein ActA which hijacks host actin machinery to propel itself into adjacent cells where it must again express LLO and a pair of phospholipases, PlcA and PlcB, for cytosolic access, thus reinitiating its infection cycle^11^. The ability of *L. monocytogenes* to adapt to life within the cytosol is achieved through multiple mechanisms including, but not limited to, modulation of specific metabolic processes, tight regulation of virulence factors to maintain the integrity of its intracellular niche, and avoidance of immune detection and host defense mechanisms^12–14^.

Menaquinone (MK) biosynthesis is an integral metabolic pathway and intermediate metabolite(s) in this pathway are required for *L. monocytogenes* cytosolic survival, independent of their known role in the production of MK^15,16^. MK acts as the sole lipid mediator of electron transport in *L. monocytogenes* and is required for a functional electron transport chain (ETC)^17^. MK biosynthesis begins with the metabolite chorismate, generated through the shikimate pathway, and is processed through a series of concurrent enzymatic reactions leading to the production of 1,4-dihydroxy-2-naphthoyl-coenzyme A (DHNA-CoA), produced by the enzyme MenB. DHNA-CoA is then converted to 1,4-dihydroxy-2-naphthoate (DHNA) by an unknown thioesterase. Lastly, DHNA is prenylated and methylated by the enzymes MenA and MenG, respectively, to generate MK^17^. A previous genetic screen revealed that the activity of MenB was required for the cytosolic survival of *L. monocytogenes*, while the enzymatic activity of MenA was not^15^. The lack of an annotated DHNA-CoA thioesterase made it impossible to know whether the virulence defects of the Δ*menB* mutant relative to the Δ*menA* mutant were due to lack of DHNA or DHNA-CoA. Therefore, it is critical to identify the unknown DHNA-CoA thioesterase in *L. monocytogenes* to better understand the respiration-independent role(s) that MK-intermediates play in its survival and virulence.

Recently, the DHNA-CoA thioesterase *YdiI* was identified and characterized in *Escherichia coli*^18^. The *L. monocytogenes* 10403s genome contains a *ydiI* homolog, LMRG_02730, which also possesses the hotdog fold domain, a feature common to other acyl-CoA thioesterases ^19,20^. In this study, we generated a genetic deletion of LMRG_02730 (renamed *menI*) in *L. monocytogenes* to determine its role in MK biosynthesis. Characterization of Δ*menI* mutants, combined with biochemical analysis of purified MenI, demonstrated that LMRG_02730 encodes the missing DHNA-CoA thioesterase. Furthermore, direct comparison between Δ*menB*, Δ*menI*, and Δ*menA* mutants allowed us to define the differential functions of DHNA-CoA, DHNA, or MK *in vitro*, *ex vivo*, and *in vivo*. Taken together, we have identified the missing DHNA-CoA thioesterase in the MK biosynthesis pathway in *L. monocytogenes* and revealed an essential role for *menI* in the cytosolic survival and pathogenesis of *L. monocytogenes*.

## RESULTS

### *Lmo2385* (*menI*) is a putative DHNA-CoA thioesterase required for menaquinone biosynthesis

Previous reports have highlighted the essentiality of MK biosynthetic intermediates in the survival and virulence of *L. monocytogenes*; however, the lack of a fully annotated pathway (**Fig. 1A**) has prevented the identification of the metabolite(s) responsible for the observed phenotypes^15,16^. Recently, the final unknown gene in the MK pathway, encoding a DHNA-CoA thioesterase, has been identified in the Gram-negative organism *Escherichia coli*, referred to as *ydiI*^18^. Sequence alignment analysis revealed that the *L. monocytogenes* 10403S genome contained a putative *ydiI* homolog, LMRG_02730 (**Fig. 1B**). Notably, both sequences contain a hotdog fold domain, a feature common to superfamily II thioesterases^19,20^. To determine the role of LMRG_02730 in *L. monocytogenes* we generated an unmarked genetic deletion mutant using allelic exchange as previously described^21^. As production of MK is essential for aerobic respiration to generate a functional membrane potential^15,17^, we hypothesized that deletion of LMRG_02730 would result in an ablated membrane potential. Wild-type *L. monocytogenes* generated a robust membrane potential that is uncoupled by the the proton ionophore CCCP. As previously reported, MK-deficient mutants Δ*menB* and Δ*menA* were unable to generate a membrane potential due to their inability to aerobically respire^15^. Consistent with its putative function as a DHNA-CoA thioeseterase, mutants lacking LMRG_02730 similarly lacked a measurable membrane potential. Membrane potential was restored for all mutants tested via either biochemical (addition of exogenous MK) or genetic complementation (**Fig. 1C**).

**Figure 1.**
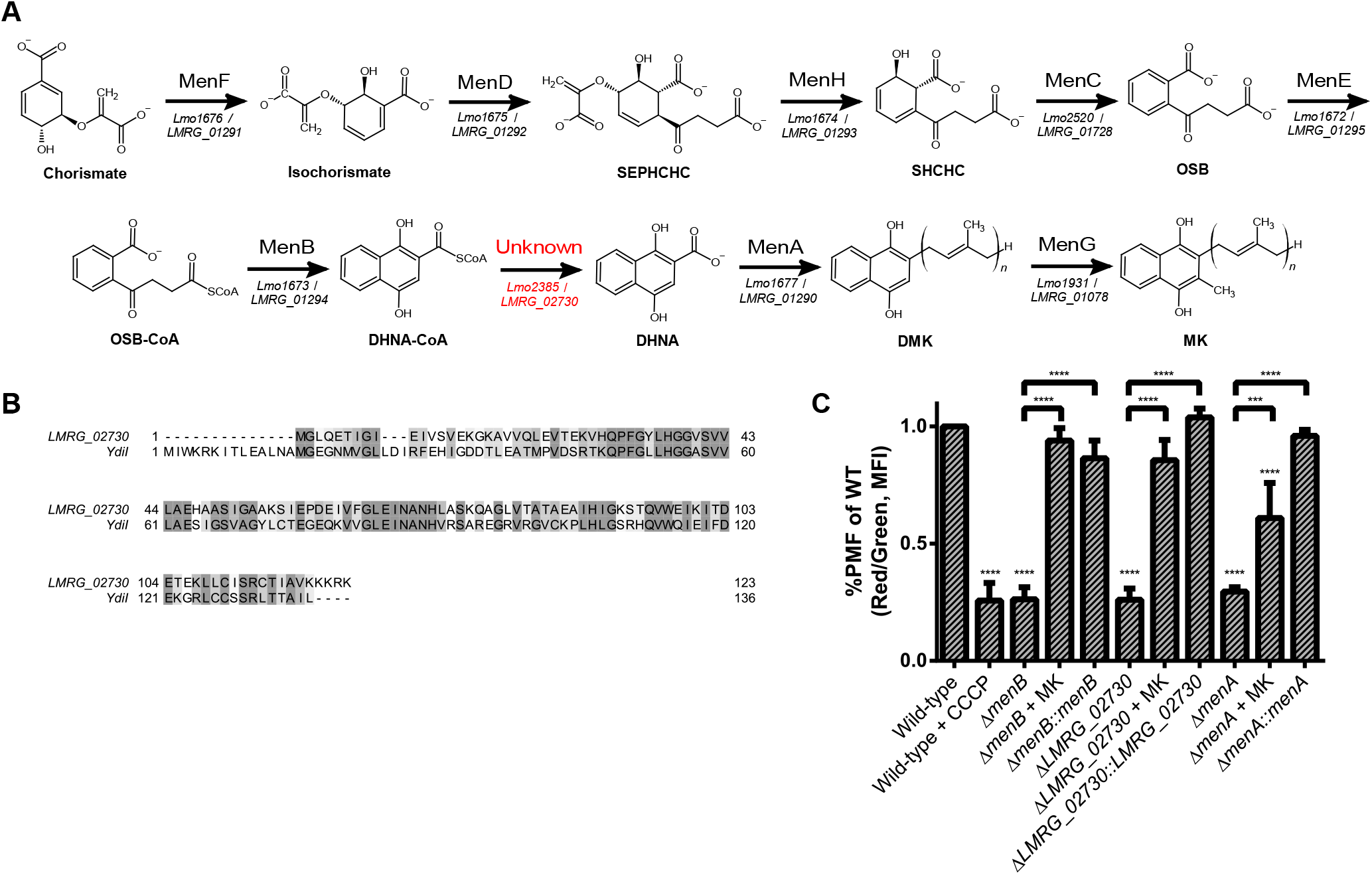
*LMRG_02730* (*Lmo2385, menI*) is homologous to *ydiI* and is required for menaquinone biosynthesis. (A) Menaquinone biosynthetic pathway in *Listeria monocytogenes*. Corresponding gene locus numbers for *L. monocytogenes* strains EGD-e (*Lmo*) and 10403S (*LMRG*; used in this study) are listed underneath reaction arrows. SEPHCHC, (1*R*,2*S*,5*S*,6*S*)-2-succinyl-5-enolpyruvyl-6-hydroxy-3-cyclohexene-1-carboxylate; SHCHC, (1*R*,6*R*)-2-succinyl-6-hydroxy-2,4-cyclohexadiene-1-carboxylate; OSB, *o*-succinylbenzoate; DMK, demethylmenaquinone. (B) Sequence alignment comparing the *Escherichia coli* protein YdiI to the *L. monocytogenes* homolog. Alignment was generated using Clustal Omega and edited using Jalview. (C) The indicated *L. monocytogenes* strains were grown in aerated BHI cultures at 37°C until mid-late logarithmic phase and then examined for membrane potentials using flow cytometry. Data represent the standard deviation of the means of three biological replicates normalized to the wild-type level (100%). Where indicated, cultures were supplemented with 5 μM MK.

To further assess whether LMRG_02730 encodes a functional DHNA-CoA thioesterase we expressed and purified the protein to perform enzyme kinetics analysis. LMRG_02730, similar to its YdiI counterpart, possesses specific hydrolytic activity towards DHNA-CoA resulting in the production of DHNA and free Coenzyme A^18^. LRMG_02730 is an active DHNA-CoA thioesterase, with a *K_m_* of 14.2 ± 8.8 μM, a *k_cat_* of 14.9 ± 3.1 s^−1^, and a second-ordered rate constant of 1.05 ± 0.08 × 10^6^ M^−1^ ∙ s^−1^. Taken together, the lack of a functional membrane potential in ΔLMRG_02730 mutants coupled with its high DHNA-CoA hydrolase activity suggests that LMRG_02730 is likely the putative DHNA-CoA thioesterase involved in MK synthesis in *L. monocytogenes*. Based on these observations we have renamed LMRG_02730 MenI.

### MenI is important for aerobic growth *in vitro* and synthesis of DHNA-CoA is essential in minimal defined media

To further evaluate the role of MenI in aerobic respiration, we compared *in vitro* growth of the Δ*menI* mutant to Δ*menB* and Δ*menA,* mutants deficient in the proteins directly upstream and downstream of MenI in MK synthesis. Under aerobic conditions in brain heart infusion (BHI) media, growth of Δ*menI*, Δ*menB*, and Δ*menA* were all similarly impaired compared to the wild-type strain and their complements (**Fig. 2A**). In contrast, under anaerobic conditions, all strains grew to similar levels as wild-type as predicted (**Fig. 2B**). These data demonstrate that the *in vitro* growth kinetics of Δ*menI* mutants phenocopies other mutants in the MK synthesis pathway, and that these defects are due to their inability to aerobically respire.

**Figure 2.**
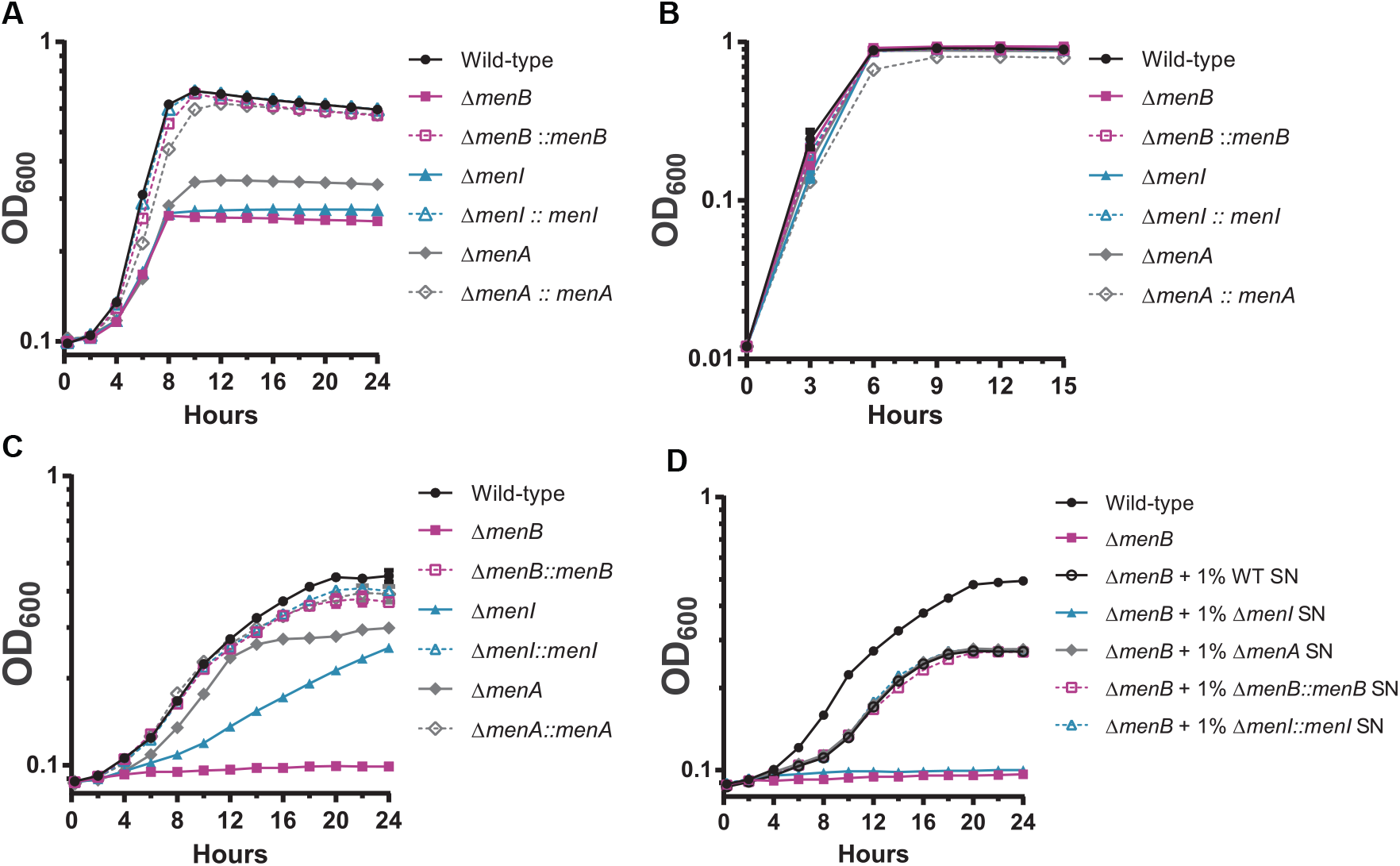
MenI is important for aerobic growth *in vitro* and synthesis of DHNA-CoA is essential in minimal defined media. *L. monocytogenes* strains, as indicated, were grown in brain heart infusion (BHI) media aerobically (A) and anaerobically (B) at 37°C. The OD_600_ was monitored for 15 to 24 h. (C) The indicated *L. monocytogenes* strains were grown in minimal defined medium at 37°C and monitored for growth (OD_600_) over 24 h. (D) Wild-type or the Δ*menB* mutant were grown in minimal defined medium at 37°C and monitored for growth (OD_600_) over 24 h. Where indicated, cultures were supplemented with 1% concentration of cell-free supernatants (SN) from wild-type, Δ*menI*, or Δ*menA* cultures grown in minimal medium. Data represent one of three biological replicates for all experiments.

In contrast to the phenotypes observed in rich BHI media, Δ*menB* mutants fail to replicate entirely in minimal defined media, whereas Δ*menA* mutants are able to grow, suggesting that BHI media contains one or multiple factors necessary to bypass DHNA-CoA or DHNA deficiency^16^. To determine whether DHNA-CoA or DHNA are required for growth in defined minimal media, we compared the growth of Δ*menI*, Δ*menB*, and Δ*menA* mutants. As previously reported, Δ*menB* mutants were incapable of growing while Δ*menA* mutants grew similar to wild-type, albeit not to the same final density (**Fig. 2C**). Δ*menI* mutants displayed an intermediate phenotype such that they grew to a similar final density as Δ*menA* mutants but at a significantly decreased rate. These data suggest that although DHNA-CoA is sufficient to promote growth in defined minimal media, the ability to produce DHNA provides an additional fitness advantage (**Fig. 2C**).

DHNA can be secreted by *Lactobacillus* spp. and *Propionibacterium* spp. and can be scavenged by *Bifidobacterium* spp. and *Streptococcus* spp. as a shared resource in complex communities^22–24^. *L. monocytogenes* can both secrete and scavenge DHNA^16^. Consistent with previous reports, cell-free supernatants from aerobically grown cultures of wild-type or Δ*menA* mutants in defined minimal media were able to rescue the growth of Δ*menB* in minimal defined media (**Fig. 2D**). In contrast, although Δ*menI* mutants can grow in minimal defined media, cell-free supernatants of Δ*menI* mutants are unable to rescue growth of Δ*menB* mutants under these conditions consistent with their inability to synthesize DHNA (**Fig. 2D**). Furthermore, these data suggest that the hydrolysis of DHNA-CoA to form DHNA is required for its subsequent secretion. Taken together, these data confirm the role of MenI for aerobic growth and DHNA secretion and further begin to define differential requirements for DHNA-CoA or DHNA under different growth conditions.

### DHNA is necessary for cytosolic survival in macrophages *ex vivo*

The activity of MenB is required for the cytosolic survival and virulence of *L. monocytogenes* in murine bone-marrow derived macrophages (BMDM) whereas the activity of MenA is dispensable^15^. To determine whether DHNA-CoA or DHNA is necessary for cytosolic survival of *L. monocytogenes*, we infected BMDMs with Δ*menB*, Δ*menI*, and Δ*menA* mutants carrying a bacteriolysis reporter plasmid to assess the levels of cytosolic killing^15,25^. *L. monocytogenes* mutants deficient in LLO (Δ*hly*) are unable to escape the host phagosome and thus act as a negative control. As previously reported^15^, Δ*menB* mutants were killed in the cytosol approximately 4- to 6-fold more frequently than the wild-type strain, while Δ*menA* mutants did not lyse under the same conditions (**Fig. 3**). Infection with Δ*menI* mutants led to levels of bacteriolysis similar to that seen in the Δ*menB* mutant (**Fig. 3**). Cytosolic survival of both Δ*menB* and Δ*menI* mutants was restored to wild-type levels upon genetic complementation. These data suggest that the synthesis of DHNA is essential for the survival of *L. monocytogenes* in macrophages.

**Figure 3.**
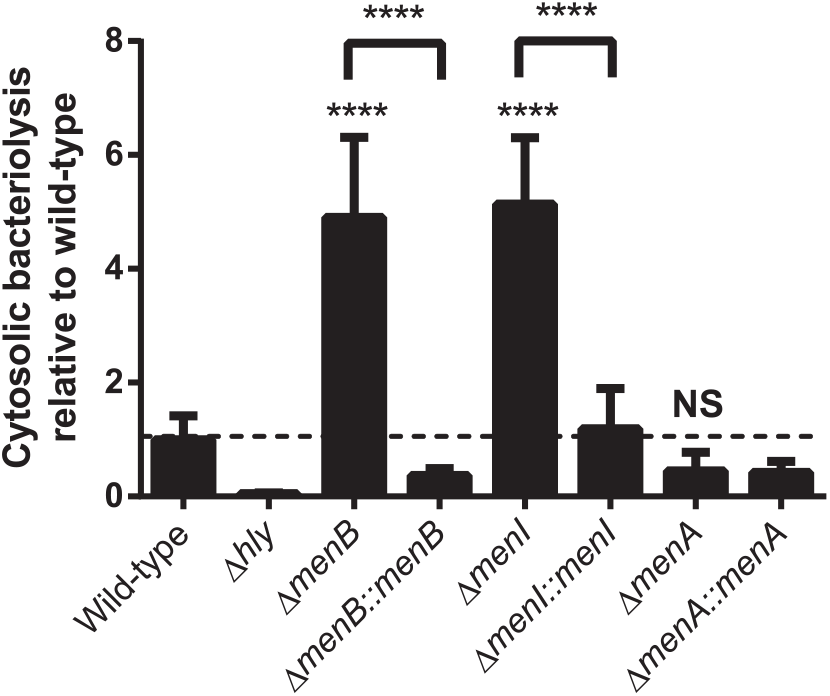
DHNA is necessary for cytosolic survival in macrophages *ex vivo*. *L. monocytogenes* MK-deficient mutants and their complements (MOI of 10) were tested for cytosolic survival in immortalized *Ifnar* ^−/−^ macrophages over a 6-h infection. Data are normalized to wild-type levels of bacteriolysis and presented as the standard deviation of the means from four independent experiments. NS, not significant.

### DHNA-CoA but not DHNA is necessary for intracellular replication and plaque formation *ex vivo*

Although DHNA is required to protect bacteria from killing in the macrophage cytosol, only a subset of bacteria ultimately undergo bacteriolysis. To further define the relative requirements for DHNA-CoA or DHNA in the *ex vivo* virulence of *L. monocytogenes*, we assessed intracellular growth of Δ*menB*, Δ*menI* and Δ*menA* mutants in primary bone marrow-derived macrophages. As previously reported^15,28^, Δ*menB* mutants are unable to replicate throughout the course of the infection (**Fig. 4A**). In contrast, both the Δ*menI* and Δ*menA* mutants were able to replicate in the macrophage cytosol, albeit not to the levels seen during wild-type infection (**Fig. 4A**). These data suggest that while DHNA is required to promote optimal cytosolic survival *L. monocytogenes* in the macrophage cytosol, only DHNA-CoA is required for bacterial replication in this environment.

**Figure 4.**
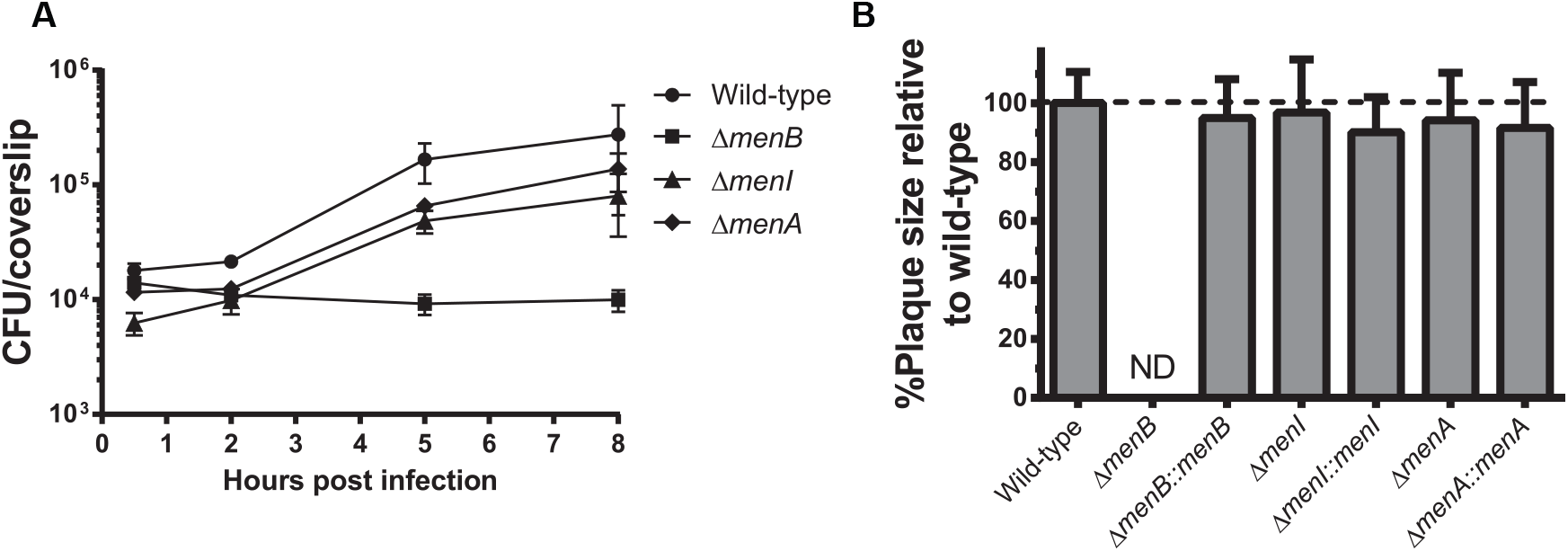
DHNA-CoA but not DHNA is necessary for intracellular replication and plaque formation in fibroblasts *ex vivo*. (A) Intracellular growth of indicated *L. monocytogenes* strains was determined in BMDMs following infection at an MOI of 0.2. Growth curves are representative of at least three independent experiments. Error bars represent the standard deviation of the means of technical triplicates within the representative experiment. (B) L2 fibroblasts were infected with indicated *L. monocytogenes* strains (MOI of 0.5) and were examined for plaque formation 4 days postinfection. Data are normalized to wild-type plaque size and represent the standard deviation of the means from three independent experiments. ND, not detected.

To further assess the relative contributions of DHNA-CoA and DHNA to *L. monocytogenes* infection, we assessed each mutant’s ability to form plaques in a monolayer of murine L2 fibroblasts^26^. To form plaques, *L. monocytogenes* must invade cells, escape from the phagosome, replicate in the fibroblast cytosol, and ultimately spread to neighboring cells using ActA-dependent actin-based motility. Consistent with previous data^15^, Δ*menB* mutants are unable to form any detectable plaques, whereas Δ*menA* mutants form plaque sizes comparable to that of wild-type (**Fig. 4B**). Importantly, plaque formation of the Δ*menI* mutant phenocopies the plaque size seen with Δ*menA* infection (**Fig. 4B**). Taken together, these data demonstrate that both DHNA and MK are dispensable for intracellular bacterial replication in macrophages and plaquing in fibroblasts. In contrast, Δ*menB* mutants are unable to replicate in macrophages and do not form plaques in a fibroblast monolayers demonstrating that DHNA-CoA is essential for intracellular replication and plaque formation.

### Deficiency in DHNA-CoA or DHNA results in organ-specific phenotypes *in vivo*

Previous reports have demonstrated that mutants deficient in MK synthesis are highly attenuated *in vivo* in a murine model of listeriosis^15,16,27,28^. Importantly, Δ*menB* mutants lacking DHNA-CoA and DHNA are significantly more attenuated than Δ*menA* mutants lacking only MK and aerobic respiration^16^. As defined above, DHNA is required for cytosolic survival in macrophages (**Fig. 3**) but is dispensable for cytosolic replication (**Fig. 4**). Therefore, to determine if DHNA and cytosolic survival contribute to virulence *in vivo* we utilized an acute murine listeriosis infection model to assess virulence of Δ*menB*, Δ*menI*, and Δ*menA* mutants and their genetic complements. As previously reported, Δ*menB* mutants were ~10,000-100,000-fold more attenuated than wild type bacteria and ~100-fold more attenuated than Δ*menA* mutants in both the spleen and liver (**Fig. 5**). Infection with Δ*menI* mutants demonstrated a tissue specific requirement for DHNA; in the spleen Δ*menI* mutants phenocopied burdens seen with Δ*menA* suggesting that in the spleen DHNA synthesis is only necessary to facilitate MK mediated respiration. In contrast, in the liver Δ*menI* mutants phenocopied Δ*menB* mutants such that they were 100,000-fold attenuated relative to wild-type bacteria, whereas Δ*menA* mutants were ~1,000-fold attenuated, suggesting a specific role for DHNA in colonization, replication, and/or survival of *L. monocytogenes* in this tissue (**Fig. 5**). The attenuated bacterial burdens of Δ*menB*, Δ*menI*, and Δ*menA* mutants *in vivo* all returned to wild-type levels upon genetic complementation (**Fig. 5**). These data suggest that DHNA, independent of respiration, is required for colonization and/or replication in the liver during disseminated *L. monocytogenes* infection. In contrast, only DHNA-CoA is necessary for colonization of the spleen in a respiration independent manner.

**Figure 5.**
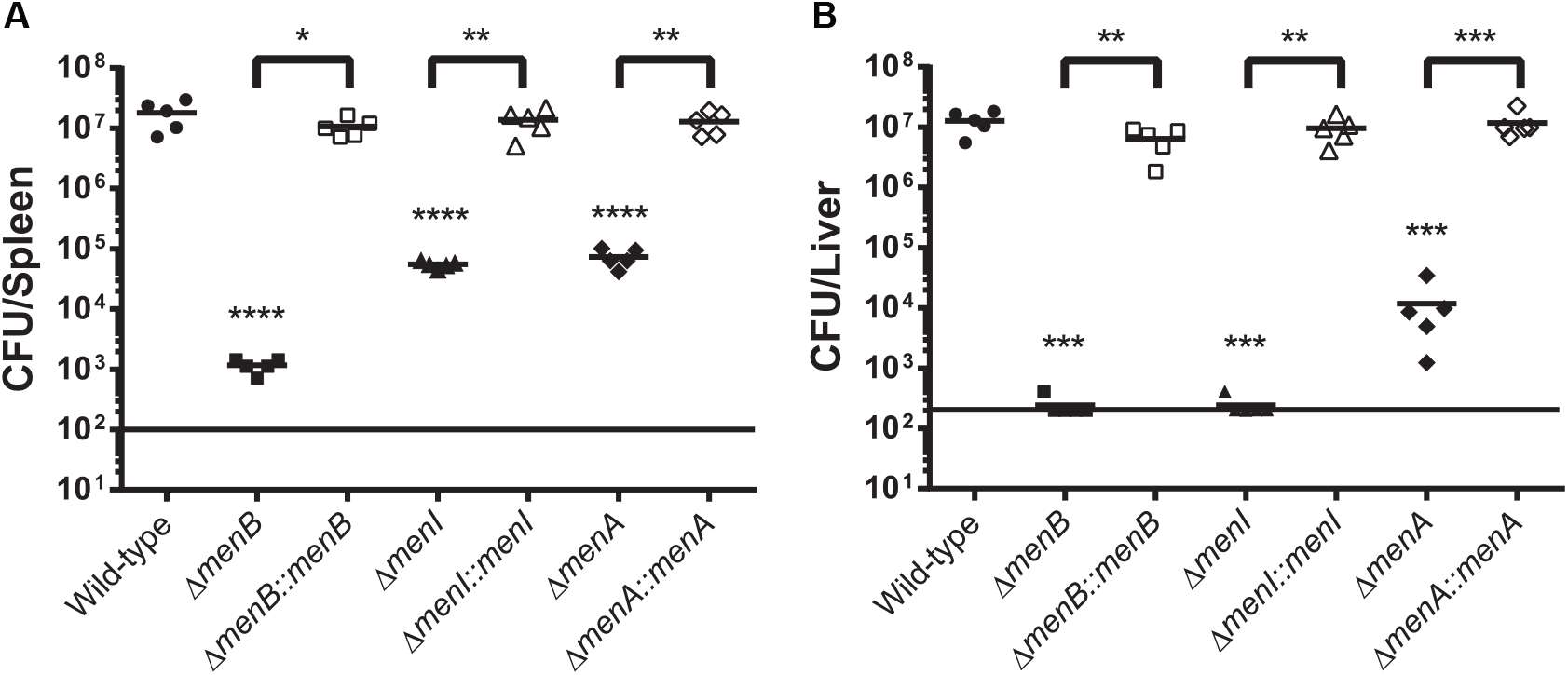
Deficiency in DHNA-CoA or DHNA results in organ-specific phenotypes *in vivo*. Bacterial burdens from the spleen (A) and liver (B) were enumerated at 48 h post-intravenous infection with 1 × 10^5^ bacteria. Data are representative of results from two separate experiments. Horizontal bars represent the limits of detection.

## DISCUSSION

Recently there has been a growing appreciation for the role of the MK biosynthesis pathway in the survival and virulence of *L. monocytogenes*^15,16,28,29^. Intermediates in the MK pathway have been shown to be essential for *L. monocytogenes* virulence, independent of their role in the production of MK and aerobic respiration^15,16^. Until now, determining whether DHNA-CoA or DHNA is the required metabolite has not been possible due to a missing gene in this pathway, a DHNA-CoA thioesterase. Here, we identified the missing gene in this pathway for *L. monocytogenes*, LMRG_02730. Using genetic and biochemical approaches, we confirmed that LMRG_02730 encodes a DHNA-CoA thioesterase, and as such we have renamed it *menI*. Additionally, infection models both *ex vivo* and *in vivo* revealed differential requirements for the synthesis of DHNA-CoA or DHNA for the survival, replication, and virulence of *L. monocytogenes*.

The canonical role of the MK biosynthesis pathway is to generate MK, which then inserts into the bacterial membrane to facilitate electron transport from NADH dehydrogenases to cytochrome oxidases in the ETC, ultimately generating a membrane potential that can drive ATP synthesis during aerobic respiration^17^. A critical outcome of ETC activity is the regeneration of the cofactor NAD^+^ from the pool of reduced NADH produced during upstream metabolic processes^29^. DHNA-CoA is the first metabolite in the MK biosynthesis pathway with a complete naphthoquinol ring (**Fig. 1A**), and DHNA itself is known to be redox capable^30,31^. Additionally, extracellular DHNA can be scavenged by *Bifidobacterium* spp. and *Streptococcus* spp. as a shared resource in complex communities and it is also known to be utilized by *L. monocytogenes* strains deficient in DHNA-CoA to stimulate growth in minimal defined media^16,22–24^. Our data reveal that synthesis of DHNA-CoA, not DHNA, is necessary to promote growth of *L. monocytogenes* in minimal defined media under aerobic conditions (**Fig. 2C**). Similarly, DHNA-CoA, but not DHNA or MK, is required for intracellular growth and plaque formation in L2 fibroblast monolayers (**Fig. 4**). One hypothesis for why DHNA and MK could be dispensable for intracellular replication is that perhaps *L. monocytogenes* can utilize the redox-capable naphthoquinol ring of DHNA-CoA to act as a cofactor to facilitate redox reactions necessary to restore, at least partially, the intracellular homeostasis of the NAD^+^/NADH ratio. Whether this process would utilize either of the two previously identified NADH dehydrogenases (LMRG_02734 and LMRG_02183) or a yet to be defined NADH dehydrogenase is unknown. Understanding the MK-independent, intracellular functions of DHNA-CoA and its role in redox balance will help illuminate novel metabolic pathways and mechanisms of replication of *L. monocytogenes* both *in vitro* and in the context of infection.

While the contribution of DHNA-CoA appears to be critical for bacterial replication *in vitro*, *ex vivo* and *in vivo*, DHNA plays a more specific role in promoting bacterial survival in the cytosol of macrophages (**Fig. 3**). This raises the intriguing question of whether DHNA acts intrinsically in the bacteria or through manipulation of the host. Recent reports have demonstrated that DHNA derived from *Propionibacterium* spp. can be used as an immune modulating molecule to reduce inflammation due to colitis in mice by suppressing macrophage-derived proinflammatory cytokines^32,33^. Furthermore, DHNA is a known agonist of the aryl hydrocarbon receptor (AhR) which is a regulator of both cellular metabolism and inflammatory responses^34,35^. Specifically in macrophages, AhR activation promotes expression of IL-10, a potent anti-inflammatory cytokine^36^. *L. monocytogenes* can secrete DHNA both *in vitro* and within the cytosol of host cells, where it can rescue the survival of DHNA-deficient strains of *L. monocytogenes* during co-infection^16^. Our data suggest that the enzymatic conversion of DHNA-CoA to DHNA is required for its subsequent secretion (**Fig. 2D**) and that synthesis of DHNA, not DHNA-CoA, is required for the cytosolic survival of *L. monocytogenes* in macrophages. How DHNA is secreted from *L. monocytogenes* and whether the bacteria intentionally secrete DHNA to act as an agonist of AhR to modulate cellular immunity remains to be determined.

Interestingly, our data also revealed organ-specific requirements for DHNA-CoA and DHNA (**Fig. 5**). Synthesis of DHNA is especially critical for the survival of *L. monocytogenes* in the liver of mice (**Fig. 5B**) however the reasons for this are currently unclear. The liver plays a key role in sterilizing and detoxifying the bloodstream, wherein over 60% of total bacteria administered via intravenous injection in an animal model is removed from the bloodstream and trapped in the liver 10 minutes post-injection^37^. Kupffer cells are liver-resident macrophages and are among the first line of defense against blood-borne bacteria, displaying a pronounced endocytic and phagocytic capacity for clearing bacteria^37,38^. Indeed, *L. monocytogenes* are ingested by Kupffer cells during infection where they induce the necroptotic death of the macrophages, leading to monocyte recruitment and an anti-bacterial type 1 inflammatory response^39^. It is possible that liver Kupffer cells or the recruited inflammatory monocytes represent a more inhospitable environment compared to the spleen that requires the synthesis and secretion of DHNA to modulate the local immune response and prevent intracellular bacteriolysis to promote the survival of *L. monocytogenes*. Future studies focused on assessing the activity of cytosolic host defenses in different myeloid cell subsets and how DHNA might specifically impact these cells could help to illuminate novel cell autonomous defense pathways. Understanding the interplay between DHNA secretion and its known role in modulating inflammatory processes will be critical to our understanding of how DHNA synthesis promotes cytosolic survival and virulence of *L. monocytogenes* in a tissue specific manner.

In summary, we have identified the unknown DHNA-CoA thioesterase in *L. monocytogenes* and have provided further evidence for the key role that the MK biosynthetic intermediates play in the survival and virulence of *L. monocytogenes*, independent of their known role in aerobic respiration. It remains to be determined whether the essential functions of DHNA-CoA or DHNA *in vitro* or during infection are correlated to basic bacterial cell physiology and/or to direct modulation of host responses. The MK biosynthesis pathway is widely conserved across the microbial kingdom, and as such identifying the underlying mechanism(s) that metabolites in this pathway play with respect to both bacterial physiology and host-pathogen interactions is critical to the development of novel therapeutic interventions.

## MATERIALS AND METHODS

### Bacterial strains, plasmid construction, and growth conditions in vitro

*L. monocytogenes* strain 10403s is referred to as the wild-type strain, and all other strains used in this study are isogenic derivatives of this parental strain. Vectors were conjugated into *L. monocytogenes* by *Escherichia coli* strain S17 or SM10^41^. Allelic exchange was performed to produce in-frame deletions of genes in *L. monocytogenes* using the suicide plasmid pksv7-oriT as previously described^21,42^. The integrative vector pIMK2 was used for constitutive expression of *L. monocytogenes* genes for complementation^43^.

*L. monocytogenes* strains were grown at 37°C or 30°C in brain heart infusion (BHI) medium (237500; VWR) or minimal medium supplemented with glucose as the sole carbon source. Minimal medium is identical to minimal medium described by Whiteley *et al*^44^. *Escherichia coli* strains were grown in Luria-Bertani (LB) broth at 37°C. For anaerobic medium studies, BHI was autoclaved and stored in an anaerobic chamber (Coy Laboratory Products) at least 24 hr before use with a gas mixture of 5% hydrogen, 20% carbon dioxide, and 75% nitrogen. *L. monocytogenes* strains were then cultured in 10 mL of conditioned BHI medium in anaerobic Hungate tubes at 37°C. Antibiotics were used at concentrations of 100 μg/ml carbenicillin (IB02020; IBI Scientific), 10 μg/ml chloramphenicol (190321; MP Biomedicals), 2 μg/ml erythromycin (227330050; Acros Organics), or 30 μg/ml kanamycin (BP906-5; Fisher Scientific) when appropriate. Medium, where indicated, was supplemented with 5 μM 1,4-dihydroxy-2-naphthoate (DHNA) (281255; Sigma) or 5 μM menaquinone (MK) (V9378; Sigma).

### Measuring bacterial membrane potential

*L. monocytogenes* strains were grown in 3 mL cultures in BHI medium at 37°C with shaking overnight. The overnight cultures were then diluted 1:100 (25 mL final volume) in 125 mL baffled flasks to mid-late logarithmic phase (OD_600_ 0.4 - 0.6) in BHI medium at 37°C, shaking at 250 rpm. Cultures were diluted to a final concentration of 1 × 10^6^ CFU/mL in sterile 1X PBS and were stained in the dark for membrane potentials using 30 μM membrane potential indicator dye DIO2(3) (3,3=-diethyloxacarbocyanine iodide) (320684; Sigma) for 30 min. The proton ionophore carbonyl cyanide 3-chlorophenylhydrazone (CCCP) (C2759; Sigma) was used to depolarize the bacterial membranes to act as a negative control and was added at a final concentration of 5 μM to control tubes prior to incubation with the membrane potential dye DIO2(3). Samples were analyzed on a BD LSR-II flow cytometer as previously described in Chen *et al*^15^.

### Synthesis of DHNA-CoA

DHNA-CoA was synthesized as previously described^45^. Briefly, 1,4-dihydroxy-2-naphthoate (102 mg, 0.5 mmol, 1 equiv) was first activated with N-hydroxysuccinimide (NHS-OH, 63.3 mg, 0.55 mmol, 1.1 equiv) in the presence of N,N’-dicyclohexylcarbodiimide (113 mg, 0.55 mmol, 1.1 equiv) in freshly distilled tetrahydrofuran (THF, 2.5 ml) for 18 hr at room temperature under magnetic stirring. After filtration and evaporation at reduced pressure, the activated product was further purified by flash column chromatography (5% methanol in dichloromethane) to obtain a brownish viscous solution (55.1 mg, 36.6%). ^1^H-NMR of the purified DHNA-NHS ester (400 MHz, chloroform-d): δ 10.28 (s, 1H), 8.30-8.16 (m, 2H), 7.74-7.66 (m, 2H), 7.11 (s, 1H), 2.83-2.73 (m, 4H).

Subsequently, DHNA-NHS (32 mg, 0.106 mmol, 4.07 equiv) was dissolved in THF (1.5 mL) and added dropwise to an aqueous solution of coenzyme A (20 mg, 0.026 mmol, 1 equiv) adjusted to pH 8.0 by sodium bicarbonate in 1.5 mL over a period of 1 hr at room temperature. Finally, the reaction was acidified by trifluoroacetic acid to pH 2.0 and extracted thrice with 10 ml dichloromethane. The aqueous solution was then purified by reversed phase high pressure liquid chromatography (HPLC) and lyophilized to yield DHNA-CoA as a brown powder (25.6 mg, 25.2%).

### MenI expression, purification, and activity assay

To express the protein, an overnight starter culture was prepared from a single colony of *E. coli* BL21 harboring the pET20b-*menI* plasmid containing a hexahistidine-tag at the N-terminus in 4 mL LB containing 50 μg/mL ampicillin. The overnight culture was used to inoculate a 1 L culture in the same media at 37°C. When OD_600_ reached 0.6, isopropyl β-D-1-thiogalactopyranoside (IPTG, 0.2 mM) was added to induce expression of MenI at 25 °C for 4 hr. The cells were harvested, resuspended in 50 mM Tris.HCl buffer containing 500 mM NaCl and 20 mM imidazole at pH 8.0, and lysed by sonication. Crude extract was then prepared and subjected to purification by nickel-nitrilotriacetic acid (Ni^2+^−NTA) metal affinity chromatography using a 5 ml HisTrap™ HP column (29-0510-21; GE Healthcare). MenI was eluted at 200 mM imidazole, desalted, and concentrated in 20 mM Tris-HCl buffer (pH 8.0) containing 10% glycerol. The purified enzyme was estimated to be >95% pure using sodium dodecyl sulfate polyacrylamide gel electrophoresis (SDS-PAGE) and its concentration was determined by UV-Vis absorbance at 280 nm using a molar extinction coefficient of 6990 M^−1^ cm^−1^ calculated from its amino acid composition with ProParam^46^. It was finally aliquoted and flash frozen in liquid nitrogen and stored at −80 °C until use.

The DHNA-CoA thioesterase activity of MenI was determined using a reported method^18^ with modifications. The extinction coefficient constants of DHNA-CoA and DHNA were measured at 396 nm and their difference was revised to be 3911 cm^−1^·M^−1^. The initial velocity was measured in triplicate at room temperature (23°C) by real time monitoring of the absorbance change at 396 nm on a Shimadzu UV-1800 UV/Visible scanning spectrophotometer in 200 mM sodium phosphate buffer (pH 8.0). The concentration of MenI was set at 13.4 nM and that of DHNA-CoA was varied from 1.8 to 57.8 μM. Km and k_cat_ were determined using Lineweaver-Burk plot.

### Transcomplementation Assays

*L. monocytogenes* strains were grown in 3 mL cultures overnight in BHI medium at 30°C without shaking overnight. The starter cultures were then diluted 1:100 into minimal media and these were then grown overnight at 37°C with shaking. Overnight aerobic cultures of the indicated strains grown in minimal medium were then centrifuged to remove bacteria. Supernatants were filtered through a 0.2 μm-pore-size syringe filter (09-740-113; Fisher Scientific). Filtered supernatants were added to wells containing indicated strains of *L. monocytogenes* in minimal media at a final concentration of 1% (vol/vol). The OD_600_ was then monitored for 24 hr (Eon, BioTek; Winooski, VT).

### Intracellular bacteriolysis assay

Standard intracellular bacteriolysis assays were performed as previously described^15^. Briefly, immortalized bone marrow-derived *Ifnar* ^−/−^ macrophages (5 × 10^5^ per well of 24-well plates) were grown in a monolayer overnight in 500 μL volume. *L. monocytogenes* strains carrying the bacteriolysis reporter pBHE573^25^ were grown at 30°C without shaking overnight. Cultures were then diluted to a final concentration of 5 × 10^8^ CFU/mL in PBS and used to infect macrophages at a MOI of 10. At 1 hr postinfection, media were removed and replaced with media containing 50 μg/ml gentamicin. At 6 hr postinfection, media from the wells were aspirated and macrophages were lysed using TNT lysis buffer (20 mM Tris, 200 mM NaCl, 1% Triton [pH 8.0]). Cell lysates were transferred to opaque 96-well plates, and luciferin reagent was added and assayed for luciferase activity (Synergy HT, BioTek; Winooski, VT).

### L2 plaque assay

Plaque assays were conducted using a L2 fibroblast cell line^26^. L2 fibroblasts were seeded at 1.2 × 10^6^ per well in a 6-well dish and infected at a MOI of 0.5 to obtain approximately 30 plaques per dish. Infection was then allowed to proceed for 1 hr, after which all wells were aspirated and washed three times with pre-warmed PBS. Wells were then overlaid with 3 mL of medium containing 0.7% agarose and 50 μg/mL gentamicin (a stock of 2 x medium with 100 μg/mL gentamicin was prewarmed at 37°C and mixed with 56°C 1.4% agarose immediately before use). At 3 days postinfection, another 3 mL of the agarose-containing medium was overlaid on each well. At 4 days postinfection, the agarose plugs were removed and cells were stained with 0.3% crystal violet for 10 min and washed twice with deionized water. Stained wells were imaged and areas of plaque formation were measured on Fiji Image analysis software^47^.

### Intracellular growth assay

BMDMs were plated on coverslips at 5 × 10^6^ cells per 60mm dish and allowed to adhere overnight. BMDMs were then infected at an MOI of with their respective strain and infection proceeded for 8 hr. At 30 min postinfection, media were removed and replaced with media containing 50 μg/ml gentamicin. Total CFU were quantified at various time points as previously described^10^.

### Acute virulence assay

All techniques were reviewed and approved by the University of Wisconsin — Madison Institutional Animal Care and Use Committee (IACUC) under the protocol M02501. Female C57BL/6 mice (6 to 8 weeks of age; purchased from Charles River) were used for the purposes of this study. *L. monocytogenes* strains were grown in BHI medium at 30°C without shaking overnight. These cultures were then back-diluted the following day 1:5 into fresh BHI medium and grown at 37°C with shaking until mid-exponential phase (OD_600_ 0.4-0.6). Bacteria were diluted in PBS to a concentration of 5 × 10^5^ CFU/mL and mice were injected intravenously with 1 × 10^5^ total CFU. At 48 hr postinfection, spleens and livers were harvested and homogenized in 0.1% Nonidet P-40 in PBS. Homogenates were then plated on LB plates to enumerate CFU and quantify bacterial burdens.

### Statistical analysis

Statistical significance analysis (GraphPad Prism, version 6.0h) was determined by one-way analysis of variance (ANOVA) with a Bonferroni posttest (*, P ≤ 0.05; **, P ≤ 0.01; ***, P ≤ 0.001; ****, P ≤ 0.0001).

## ACKNOWLEDGEMENTS

We would like to thank Robert L. Kerby and the laboratory of Dr. Federico Rey for use of their anaerobic chamber as well as technical assistance with anaerobic media and anaerobic tube preparation.

This work was funded by the National Institutes of Health (T32007215 [HBS] and R01AI137070 [J-D S]). This work was also partially supported by the National Natural Science Foundation of China (21877094 [ZG]) and the Hong Kong Branch of Guangdong Southern Marine Science and Engineering Laboratory (Guangzhou) (SMSEGL20SC01-E [ZG]). The funders had no role in study design, data collection and interpretation, or the decision to submit the work for publication.

